# Inferotemporal cortex stimulation is perceived before its onset

**DOI:** 10.64898/2026.10.02.756371

**Authors:** Elia Shahbazi, Drew Nguyen, Timothy Ma, Arash Afraz

**Author notes:** These authors contributed equally to this work (Elia Shahbazi, Drew Nguyen).

## Abstract

Neural response latency measures when stimulus-evoked activity becomes detectable, but it does not reveal when the brain assigns the causal contribution of that activity to events in the external world. We define this temporal adjustment as projected delay and measure it in the macaque inferotemporal cortex using optogenetic stimulation and behavior. Two rhesus macaques performed a cortical perturbation detection task in which a visual masker appeared at different times relative to stimulation. Detection was selectively impaired when the masker began 200 ms before stimulation. We then tested this timing using Perceptography, a method that optimizes image perturbations to mimic stimulation-evoked visual events. Image perturbations presented 200 ms before stimulation elicited higher false-alarm rates than perturbations presented at stimulation onset. Here, we show that stimulation-evoked visual events in inferotemporal cortex depend preferentially on visual information originating approximately 200 ms before the cortical perturbation. This projected delay exceeds typical inferotemporal response latencies and suggests temporally extended readout mechanisms relevant to causal models of visual perception and the timing of cortical visual prostheses.

## Introduction

Neurons across the visual cortex display distinct response latencies, reflecting their position within the visual hierarchy and other circuit-level properties^1,2^. A single sensory event, therefore, activates different brain areas at different times. To support coherent perception and make the correspondence between an external event and its resulting asynchronous responses across various areas, the brain must take into account a temporal offset for each area separately and compensate for it. A neural event, then, should be projected back by the brain to a physical event that happened earlier. The amount of this temporal adjustment reflects the brain’s assumption about the functional delay of a given area. This temporal adjustment, defined here as the “projected delay”, is conceptually distinct from “response latency”, the traditional measure of neural delay.

Response latency is defined as the earliest time a neuron’s firing rate exceeds its baseline activity. This measure is correlational in nature and indicates when the neural response becomes detectable by an external observer. Nevertheless, this correlational measure does not reveal the functional delay that the brain causally assigns to a given area during its function.

Consequently, isolating the projected delay demands a causal assay rather than a correlational observation. The projected delay cannot be recovered from passive recordings alone; it requires a time-locked exogenous perturbation of the cortex and a behavioral readout of when the stimulation-induced percept is judged to have occurred relative to a reference stream of physical events. Only such a test can cleanly dissociate projected delay from sensory-driven response latency.

This framing yields three testable hypotheses about the relation between causal and correlational delays (Fig. 1c): simple decoding (causal approximately equals correlational; spikes become perceptually effective as soon as they arrive), consistent with rapid readout of inferotemporal population activity^3-5^; predictive decoding (causal < correlational; anticipatory mechanisms shift perceived time forward), consistent with predictive temporal-alignment accounts^6,7^; and processing deferral (causal > correlational; additional readout or integration is required).

**Fig 1.**
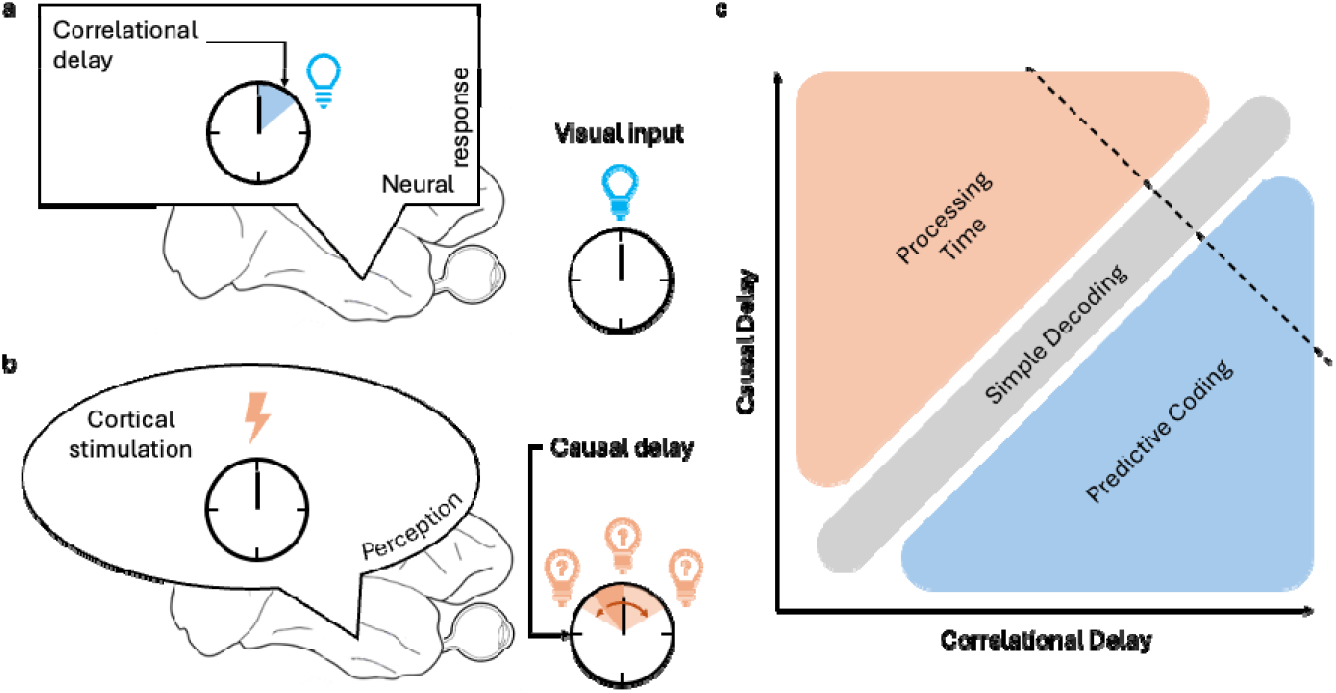
Correlational delay and projected delay. a, Correlational delay, also known as response latency, is the interval between optical presentation of a visual stimulus and emergence of its corresponding neural activity, measured by a statistically significant increase in firing rate, decoding performance or another relevant measure. b, Projected delay is defined here as the interval between an exogenous increase in neural activity, induced by cortical stimulation, and the perceived time assigned to the evoked visual event. c, Three hypotheses based on the relative durations of causal and correlational delays: predictive decoding, simple decoding and processing deferral.

**Fig 2.**
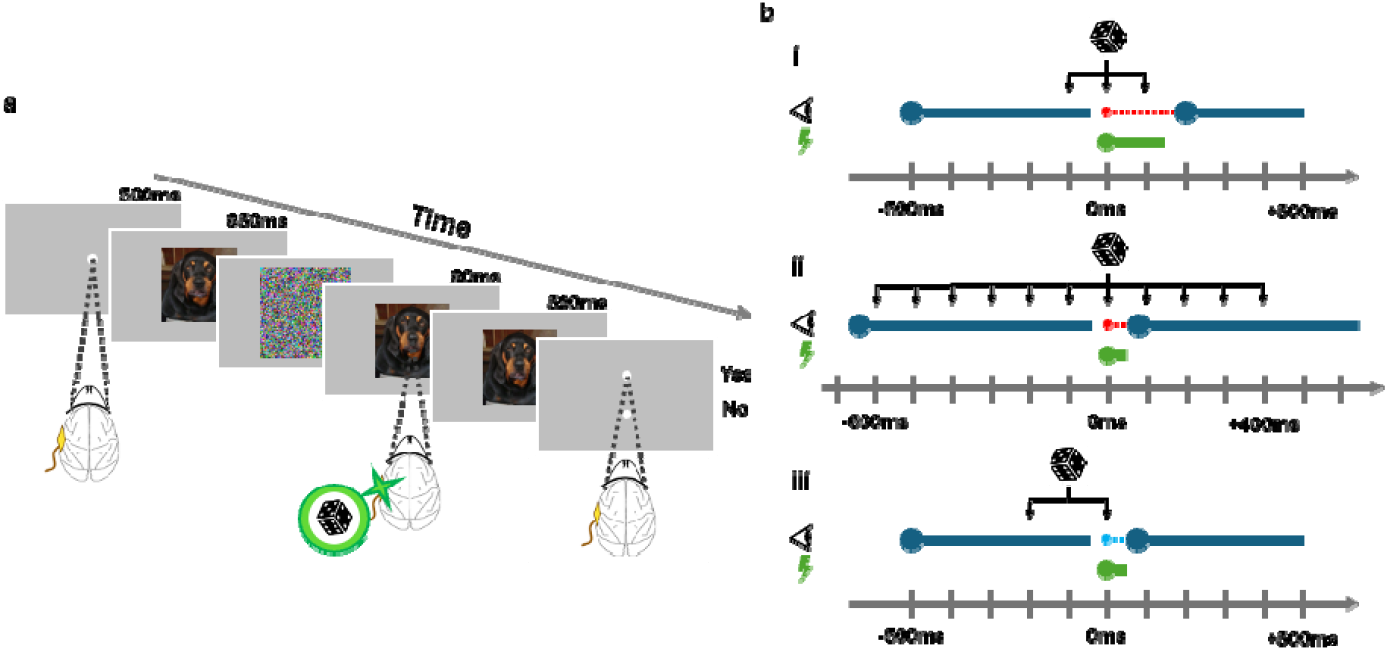
Masked cortical perturbation detection task and experimental timing. a, Task overview. The monkey maintains fixation while an image is presented; optogenetic stimulation is delivered on half of trials and a visual masker or image perturbation is presented at a controlled temporal offset. After image presentation, the monkey reports whether stimulation occurred by saccading to one of two response targets. b, Timing schemes for the narrow-range masking experiment, broad-range masking experiment and Perceptography validation experiment. Exact durations and onset times are reported in Methods.

A lack of causal understanding of projected delays poses a practical challenge for brain-stimulation experiments and attempts to restore vision by cortical stimulation: the perceived timing of a cortical stimulation event relative to an external clock remains unknown. Artificial stimulation of inferotemporal cortex can alter visual behavior and produce image-dependent detectable visual events^8,9^. This challenge becomes explicit in Perceptography^10^, which uses precisely timed image optimization to reconstruct perceptual events induced by local stimulation in high-level visual cortex. In Perceptography, a monkey fixates on a seed image that is briefly replaced by a perturbed image. In half of the trials, randomly chosen, inferotemporal cortex is optogenetically stimulated. After the image sequence, two response targets appear, and the animal reports whether brain stimulation occurred. By tracking false alarms on non-stimulated trials and using machine-learning-guided image evolution over many thousands of trials, the process optimizes image perturbations to maximize the behavioral false-alarm rate. The resulting confusion depends not only on matching what the stimulation-induced percept looks like, but also on matching when it is perceived: the on-screen perturbation and the internal percept must overlap in time in the animal’s experience.

When Perceptography was introduced in 2024^10^, the temporal position of the stimulation-induced percept on the physical timeline of the trial was unknown. To hedge against this uncertainty, we used relatively long image-perturbation and stimulation epochs, increasing the chance that the two events would overlap in perception regardless of the true projected delay. That strategy allowed Perceptography to work, but it left the temporal alignment between screen perturbation and hallucination underspecified, and it could not reveal how far back in time the brain projects the causal consequences of inferotemporal stimulation. Correlational measures such as response latency cannot fill this gap because they indicate when neural activity becomes detectable to an outside observer, not when that activity is endogenously assigned to a particular moment in the external stream of events.

A direct estimate of the projected delay is therefore required to temporally optimize Perceptography: it indicates where on the physical timeline to place a brief image perturbation so that its perceptual impact maximally overlaps the stimulation-evoked event. Knowing this offset allows us to shorten the stimulation and image-perturbation windows, maximize temporal overlap and make behavioral image optimization more precise and efficient. More generally, direct measurement of projected delay provides temporal information relevant to real-time cortical visual prostheses.

Visual masking provides a well-established way to probe the temporal dynamics of visual processing^11-14^. Here, we investigate projected delay using optogenetic stimulation in macaque inferotemporal cortex and a modified cortical perturbation detection task in which a masker appears at one of several randomly selected times relative to stimulation. We first use a coarse temporal range to identify when masking is most effective, then repeat the experiment over a broader range with finer stimulation timing. Finally, we test the inferred timing using Perceptography. Across both animals, masking most strongly reduces stimulation detection when presented approximately 200 ms before stimulation, and image perturbations presented at this earlier time produce higher false-alarm rates than perturbations presented at stimulation onset. These results indicate that stimulation-evoked visual events in inferotemporal cortex depend preferentially on visual information from an earlier temporal reference frame.

## Results

### Narrow-range masked CPD experiment

In our initial experiment, two rhesus macaques (Kr and Ph) were trained to perform a masked cortical perturbation detection (CPD) task. The macaques fixated on a central fixation point for 500 ms to initiate a trial. In each trial, an image was presented on-screen for 1000 ms; in 50% o trials, chosen randomly, optogenetic stimulation was applied to inferotemporal cortex 500 ms into image presentation for 100 ms. Following image presentation, the macaques made a saccade to one of two response targets, with a liquid reward for correct responses. In every trial, a grey square (the “masker”) appeared at one of three possible times: 100 ms before stimulation, simultaneously with stimulation or 100 ms after stimulation (-100 ms, 0 ms and +100 ms relative to stimulation, respectively). We have previously shown that macaques rely on stimulation-evoked visual events to detect inferotemporal optogenetic stimulation and that detection interacts with the image presented on the screen^9,10^. In the first two experiments here, we sought to interfere with the animals’ perception of the stimulation-evoked visual event, resulting in a decrease in task performance measured by d’ (d prime). By presenting the masker at different temporal offsets from stimulation and examining performance as a function of masker onset time, we used an external visual event as a reference for the temporal position to which the cortical stimulation event is projected. Control trials without the masker were included to validate its effectiveness. See Methods for full experimental details.

In this initial experiment, Kr completed 4928 trials and Ph completed 10222 trials. The only significant pairwise difference in d’ across masker onset times was between -100 ms and 0 ms for Kr (p < 0.05; two-tailed permutation test, d’ difference). In Ph, the comparisons between -100 ms and +100 ms and between 0 ms and +100 ms approached significance in the same direction but did not reach the threshold (p = 0.068 and p = 0.054; two-tailed permutation test, d’ difference; Fig. 3a). We also found a significant positive linear relationship between masker onset time and d’ for Ph, suggesting that detection performance decreased as the masker was presented earlier in the trial (p < 0.05, R^2^ = 0.7296; two-tailed permutation test, slope). Kr showed a positive but non-significant trend (p > 0.1, R^2^ = 0.5428; two-tailed permutation test, slope). Masker validation (Fig. 3b) showed that trials in which the masker appeared had significantly lower d’ than trials without the masker (Kr: p < 0.05; Ph: p < 0.001; one-tailed permutation test, d’ difference). These results show that the masker decreased performance and suggest a greater effect at earlier onset times. Because the trends were not conclusive across both animals, we next expanded the temporal range and increased temporal resolution.

**Fig 3.**
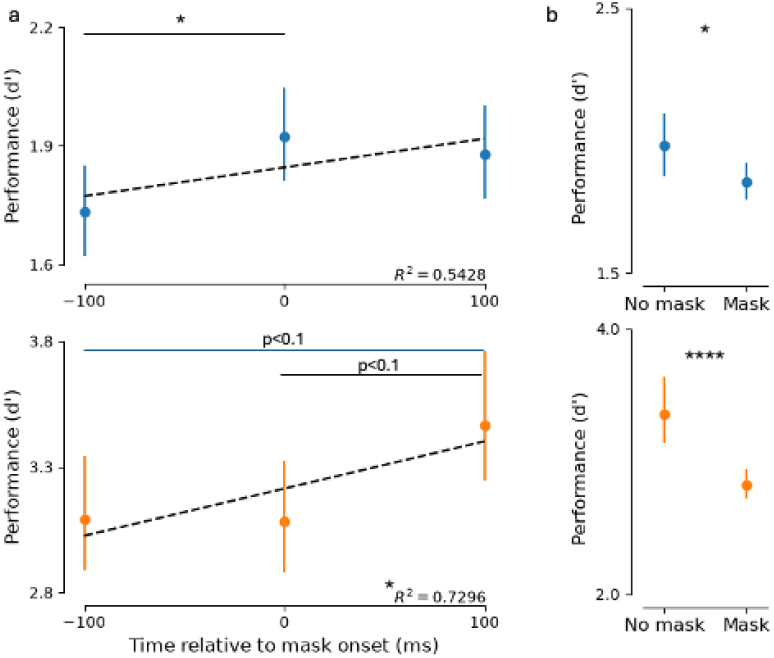
Narrow-range masked CPD performance. a, CPD performance when the masker was presented 100 ms before stimulation, at stimulation onset or 100 ms after stimulation. Kr is shown in blue and Ph in orange. Error bars show bootstrapped 95% confidence intervals. R^2^ values are from linear fits. Pairwise significance was assessed by two-tailed permutation tests. b, CPD performance on unmasked versus masked trials. Error bars show bootstrapped 95% confidence intervals. Masked and unmasked conditions were compared using one-tailed permutation tests. Kr completed 4928 trials and Ph completed 10222 trials.

### Broad-range masked CPD experiment

Although the first experiment suggested that earlier masker onset produced a greater drop in performance, the trend was not significant across both animals and did not identify the time at which masking was maximally effective. If the underlying relationship were positive, it must also have a temporal limit. We therefore repeated the masked CPD experiment in Kr and Ph using a broader range of masker onset times and shorter optogenetic stimulation. Images were presented for 1300 ms. A 100-ms masker appeared at one of 11 time points from -600 ms to +400 ms relative to stimulation, separated by 100-ms intervals. The masker was changed to a high-contrast, colorful, animated noise pattern, and its effect was again validated using trials with no masker. Optogenetic stimulation was applied on 50% of trials, randomly selected, beginning 650 ms after image onset and lasting 60 ms. See Methods for complete details.

In this experiment, Kr completed 25880 trials and Ph completed 120344 trials. The only masker onset time that elicited a significant decrease in d’ compared with all other conditions was -200 ms relative to stimulation (Fig. 4a; p < 0.01 for both Kr and Ph; one-tailed permutation test, one-versus-rest d’ contrast). Masker validation (Fig. 4b) showed that trials in which the masker appeared had significantly lower d’ than trials without the masker (p < 0.0001 for both Kr and Ph; one-tailed permutation test, d’ difference). Thus, with a broader temporal range and shorter stimulation, masking interfered most strongly with detection when presented 200 ms before stimulation, suggesting that detection of stimulation-evoked visual events depends on visual information preceding stimulation.

**Fig 4.**
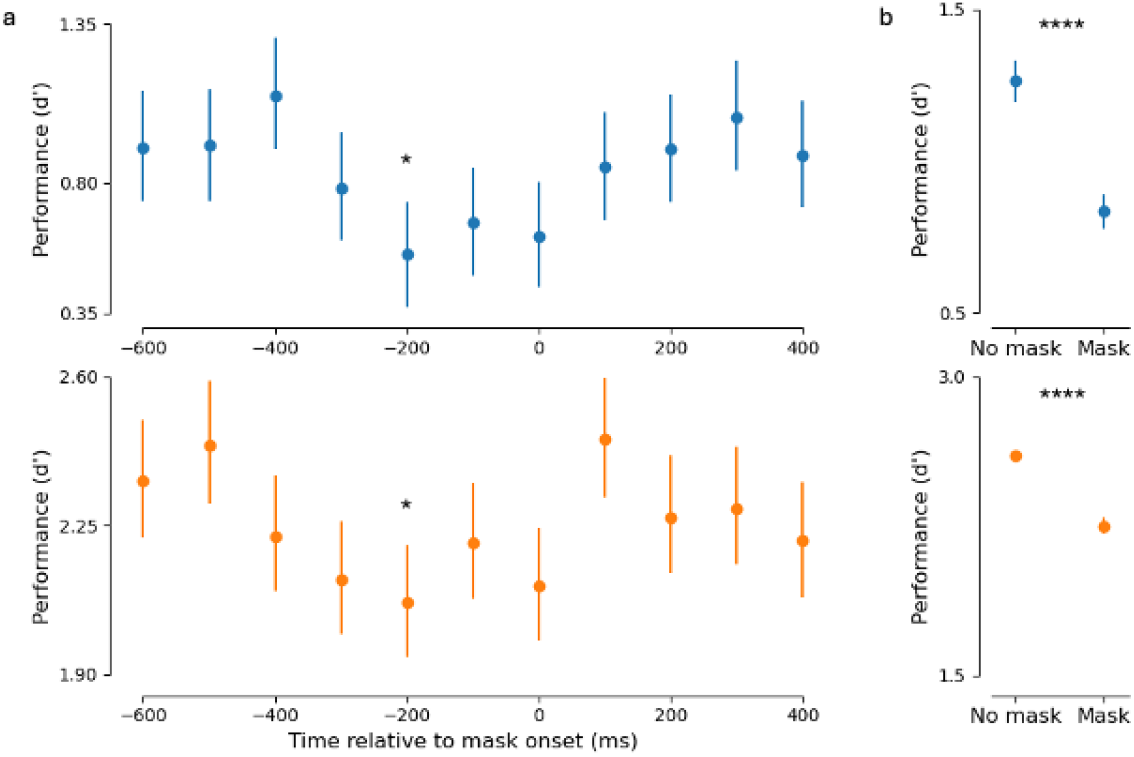
Broad-range masked CPD performance. a, CPD performance as a function of masker onset time from -600 ms to +400 ms relative to stimulation. Kr is shown in blue and Ph in orange. Error bars show bootstrapped 95% confidence intervals. For each time point, significance was assessed as the one-versus-rest contrast in d’ across 10,000 permutations; the asterisk marks p < 0.01. b, CPD performance on unmasked and masked trials. Error bars show bootstrapped 95% confidence intervals; four asterisks indicate p < 0.0001 from one-tailed permutation tests. Kr completed 25880 trials and Ph completed 120344 trials.

### Perceptographic validation

The masked CPD experiments indicated a projected delay centered approximately 200 ms before stimulation. To test this estimate in an independent behavioral perturbation paradigm, we performed Perceptography as previously described^10^, presenting image perturbations either at stimulation onset or 200 ms before stimulation. If stimulation-evoked visual events in inferotemporal cortex are assigned to an earlier temporal reference frame, image perturbations presented 200 ms before stimulation should more closely resemble the stimulation-evoked event and should therefore produce a higher false-alarm rate.

Kr completed 12120 trials and Ph completed 17229 trials. Image perturbations presented 200 ms before stimulation had significantly higher false-alarm rates than those presented at stimulation onset (Kr: p < 0.001; Ph: p < 0.05; statistical test as specified in Methods and Fig. 5). This result is consistent with the approximately 200-ms projected delay inferred from the masking experiments.

**Fig 5.**
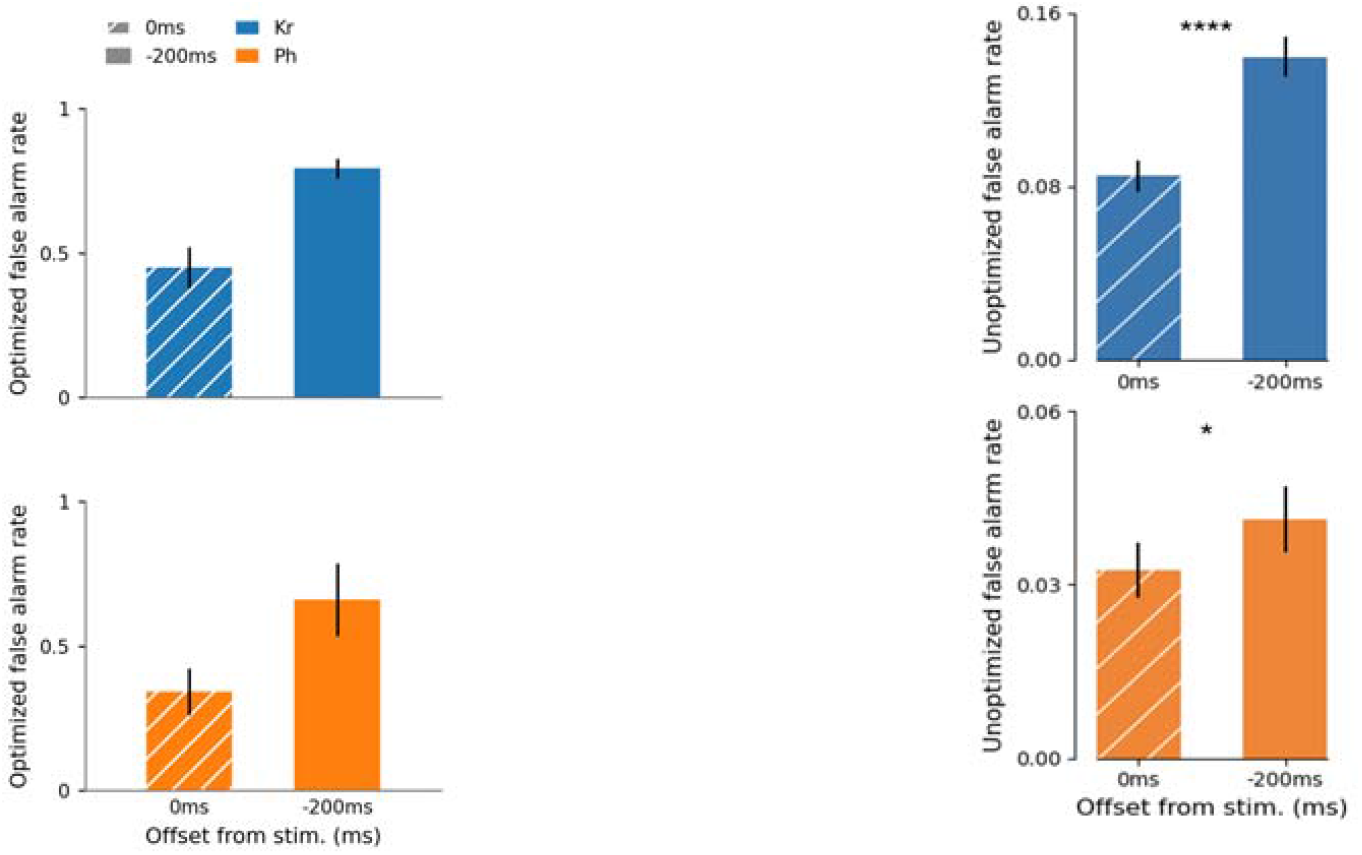
Perceptographic validation at 0 ms and -200 ms relative to stimulation. Left, optimized false-alarm rates for the best-performing image families in Kr (blue) and Ph (orange). For each timing condition, the two best-performing image families were selected based on cumulative family false-alarm rate; image families in the 0-ms condition survived two optimization rounds, whereas selected -200-ms families survived six rounds. Right, mean false-alarm rates for unoptimized, randomly generated image perturbations. Error bars show bootstrapped 95% confidence intervals. One asterisk denotes p < 0.05 and four asterisks p < 0.0001. Kr completed 12120 trials and Ph completed 17229 trials.

## Discussion

Our data show that detection of stimulation-evoked visual events is maximally hindered by visual noise presented approximately 200 ms before stimulation. Perceptography provides convergent evidence: image perturbations presented at -200 ms produce higher false-alarm rates than perturbations presented at stimulation onset. Together, these results are consistent with a projected delay of approximately 200 ms for stimulation-evoked visual events in inferotemporal cortex.

Response latency of macaque inferotemporal neurons has been extensively studied and ranges from approximately 82 to 118 ms depending on visual stimulus and analysis method^15,16^. The behavioral projected delay measured here exceeds this range and the timing expected from rapid neural decoding studies, favoring a processing-time account over instantaneous readout. The divergence between behavioral and decoder latency suggests that perception may rely on a slower, integrating readout than classifiers trained on neural activity. For image-optimization frameworks that aim to match the perceptual contribution of a neuronal event, our findings therefore suggest that the temporal relationship between the cortical event and retinal input should be explicitly measured rather than inferred from response latency. The same consideration is relevant to real-time cortical visual prostheses.

In our comparison of Perceptography with image perturbations presented at 0 ms versus -200 ms relative to stimulation, the earlier perturbations produced higher false-alarm rates. We subsequently adopted this timing in Perceptography and observed consistent gains while shortening the image-presentation sequence around the predicted temporal window. Shorter trials increase data-collection speed and expand the practical design space for future Perceptography experiments.

Libet originally proposed that conscious sensory experiences can be retrospectively referred back in time to coincide with an earlier peripheral time-marking event, based on somatosensory and thalamocortical stimulation experiments in humans^17^. The interpretation of those experiments and the broader backward-referral account have remained debated^18,19^. Our findings are consistent with a backward temporal adjustment in vision, but the present behavioral assay is indirect. The macaques do not explicitly report the order of a cortical stimulation event and a concurrent image change; instead, we infer temporal alignment from the effect of masking on stimulation detection. A direct temporal-order judgment would therefore provide a stronger test of whether the perceived order of internal and external events is dissociated from their physical order. A second limitation is that the OptoArray used here does not permit simultaneous electrophysiological recording, so the neural population dynamics associated with the approximately 200-ms behavioral effect remain unknown.

In summary, visual interference selectively impairs detection of inferotemporal optogenetic stimulation when presented approximately 200 ms before stimulation, and image perturbations mimic stimulation-evoked visual events more effectively at -200 ms than at 0 ms. These results indicate that stimulation-evoked visual events incorporate visual information originating from an earlier time point, consistent with a temporally extended readout. The observed projected delay exceeds conventional estimates based on response latency and neural decoding, raising questions about the time course and computational complexity of the brain’s native readout.

Future work combining causal perturbation, behavioral timing and neural population recording will be required to identify the mechanisms that implement this temporal alignment and to determine how projected delay depends on cortical site, attention and anticipation.

## Methods

All procedures were conducted in accordance with and approved by the National Institute of Mental Health Animal Care and Use and Committee guidelines.

### Surgical procedure and optical array implantation

We injected viruses into an approximately 5 x 5 mm area of central inferotemporal cortex in two male macaque monkeys (*Macaca mulatta*), Kr (5 years old, approx. 8 kg) and Ph (8 years old, approx. 14 kg)., to express the excitatory opsin C1V1 under the CaMKIIa promoter. We introduced AAV5-CaMKIIa-C1V1(t/t)-EYFP using a custom injection apparatus at a concentration of 8 x 10^12 particles per milliliter. The apparatus contained four 31-gauge needles arranged in a 2 x 2 mm square grid. To encompass the central region of inferotemporal cortex, we performed a sequence of four injections at sixteen sites, collectively generating a transduced area of approximately 6 x 6 mm. At each site, 10 microliters of viral solution was administered at 0.5 microliters per minute, for a total volume of 160 microliters. We waited 10 min after each injection before retracting the needles to allow diffusion into the cortex.

We then implanted an OptoArray (Blackrock Neurotech; light-emission peak 530 nm) in the virally injected regions. The devices were placed 7-12 mm anterior to the interaural line, spanning TEpd to TEad as delineated in the Saleem and Logothetis atlas. The OptoArray enables rapidly reversible optical stimulation of approximately 1 mm3 cortical subregions but does not permit neural recording. Technical details of the OptoArray and surgical protocols are described in previous reports^9,10^.

### Testing configuration and ethics

All experiments were performed in a well-illuminated chamber to prevent retinal dark adaptation and minimize the possibility that the animals could detect cortical light stimulation through cranial structures. The animals were positioned 57 cm from a 32-inch IPS LCD display (1920 x 1080 resolution, 120 Hz; Cambridge Research Systems). Data acquisition was controlled by a custom MWorks script running on a Mac Pro, and eye movements were tracked with an EyeLink 1000 Plus system (SR Research).

### Training

Kr and Ph were initially trained to detect brief optogenetic stimulation in inferotemporal cortex while fixating on images generated by a generative adversarial network in a cortical perturbation detection task. In each trial, the monkeys fixated on a central target, followed by a 1000-ms presentation of a generated image. In half of trials, chosen randomly, optogenetic stimulation was delivered to inferotemporal cortex for 150 ms beginning 500 ms after image onset. After image presentation, the monkeys saccaded to one of two response targets to report whether stimulation was delivered, with a liquid reward for correct responses.

After CPD training, each monkey was trained on the Perceptography paradigm^10^. In each trial, the monkeys fixated on a central target, followed by a 500-ms presentation of a generated seed image. The image briefly changed to a perturbed version for 150 ms before reverting to the seed image for 350 ms. In half of trials, chosen randomly, optogenetic stimulation was delivered to inferotemporal cortex. After image presentation, the monkeys reported whether stimulation had occurred by saccading to one of two response targets.

### Narrow-range masked CPD: data collection

The monkeys first performed a preliminary version of the masked CPD task. They fixated for 500 ms to initiate a trial, after which a generated image was presented for 1000 ms. A grey square with the same luminance as the monitor background appeared for 120 ms at one of three time points, obscuring the presented image. The masker appeared 100 ms before stimulation, simultaneously with stimulation or 100 ms after stimulation. In half of trials, chosen randomly, optogenetic stimulation was applied for 100 ms beginning 500 ms after image onset. After image presentation, the monkeys saccaded to one of two response targets and received a liquid reward for correct responses.

To assess masker efficacy, we also collected unmasked trials with the same structure except that no masker appeared and image presentation was uninterrupted.

### Narrow-range masked CPD: data analysis

Trials were grouped by masker onset time (-100 ms, 0 ms or +100 ms relative to stimulation). Detection performance was quantified using d’ (the difference between the Z-transforms of hit rate and false-alarm rate). We estimated 95% confidence intervals by bootstrapping 10,000 resamples. Because we did not have a directional hypothesis at this stage, pairwise differences in d’ between masker-time conditions were assessed using two-tailed permutation tests with 10,000 resamples. We also fit a linear regression to performance as a function of masker onset time.

To validate masker efficacy, we compared d’ between masked and unmasked trials. Because the masker was designed to reduce detection performance, we used a one-tailed permutation test with 10,000 resamples and estimated 95% confidence intervals by bootstrapping 10,000 resamples.

### Broad-range masked CPD: data collection

To characterize the temporal extent of the masker effect, we expanded the range of masker onset times. The monkeys again fixated for 500 ms to initiate a trial. An image was presented for 1300 ms, and a 100-ms masker appeared at one of 11 time points from -600 ms to +400 ms relative to stimulation, separated by 100-ms intervals. The masker was a full-contrast animated noise pattern composed of randomly chosen frames regenerated for each session. In half of trials, chosen randomly, inferotemporal cortex was stimulated for 60 ms. After image presentation, the monkeys reported whether stimulation occurred by saccading to one of two response targets.

We again collected unmasked control trials identical in structure except that no masker was presented.

### Broad-range masked CPD: data analysis

Trials were grouped by masker onset time and performance was quantified using d’. We estimated 95% confidence intervals by bootstrapping 10,000 resamples. For each time point, we tested whether d’ was lower than the pooled d’ of all other masker-onset conditions using a one-tailed permutation test on the contrast d’(condition) - d’(all other conditions), with 10,000 resamples. The analysis used an alpha threshold of 0.01. Masked and unmasked trials were compared using a one-tailed permutation test with 10,000 resamples, and 95% confidence intervals were estimated by bootstrapping 10,000 resamples.

### Perceptography validation: data collection

We next tested whether the temporal effect identified in masked CPD generalized to Perceptography. The monkeys fixated on a central point for 500 ms to initiate a trial. A generated seed image was presented for 1000 ms. In half of trials, chosen randomly, optogenetic stimulation was applied to inferotemporal cortex. On each trial, the seed image changed to a matched image perturbation either at stimulation onset or 150 ms before stimulation. The same image perturbations were used across the two onset conditions. Previous work showed that such perturbations can increase false-alarm reports on non-stimulated trials^10^. After image presentation, the monkeys reported whether stimulation occurred by saccading to one of two response targets^20-32^

### Perceptography validation: data analysis

We analyzed the mean false-alarm rate of unoptimized image perturbations and the behavior of optimized image families. In the current analysis description, mean false-alarm rates for unoptimized perturbations are compared between onset conditions using a two-tailed t-test. For optimized image families, the proportion of images in each pool that generated offspring in the next iteration was used as a measure of survival, and survival-rate p values were estimated by permutation testing per iteration with confidence intervals estimated by 10,000 bootstrap resamples^22,23,24,25,26,27,28,29,30,21,32^

### Statistics and reproducibility

The study included two macaque monkeys. Trial-level counts for each experiment are reported in the Results and figure legends. Stimulation assignment and masker-onset conditions were randomized within the task as described above. Confidence intervals were estimated by bootstrap resampling and inferential tests used permutation procedures except where otherwise specified.

## Data availability

The data and material that support the findings of this study will be available upon the request. Seed_image collection: https://github.com/eliashahbazi/Seed_collection.git.

## Code availability

Pretrained model that we used in DaVinci and Ahab https://tfhub.dev/deepmind/biggan-deep-128/1.

The code is available upon request. Interested parties should contact Elia Shahbazi at for access.

## Acknowledgements

We thank Mark Eldridge and Reza Azadi for their crucial contributions to the surgeries. We thank Reza Azadi for the initial training of monkey Ph. This research was supported by the Intramural Research Program of the NIMH ZIAMH002958 (to A.A.)

## Author contributions

Concept: E.S., T.M., Experiment Design: E.S., T.M., A.A., Data collection and analysis: E.S., D.N. and T.M. Writing the manuscript: E.S., A.A, D.N., and T.M.

